# Challenges in Identifying and Interpreting Organizational Modules in Morphology

**DOI:** 10.1101/097261

**Authors:** Borja Esteve-Altava

## Abstract

Form is a rich concept that agglutinates information about the proportions and topological arrangement of body parts. Modularity is readily observable in both the variation of proportions (variational modules) and the organization of topology (organizational modules). The study of variational modularity and of organizational modularity faces similar challenges regarding the identification of meaningful modules and the validation of generative processes; however, most studies in morphology focus solely on variational modularity, while organizational modularity is much less understood. A possible cause for this bias is the successful development in the last twenty years of morphometrics, and specially geometric morphometrics, to study patters of variation. This contrasts with the lack of a similar mathematical framework to deal with patterns of organization. Recently, a new mathematical framework has been proposed to study the organization of anatomical parts using tools from Network Theory, so-called anatomical network analysis. This essay explores the potential use of this new framework – and the challenges it faces in identifying and validating biologically meaningful modules in morphological systems –, by providing an example of a complete analysis of modularity of the human skull and upper limb. Finally, we suggest further directions of research that may bridge the gap between variational and organizational modularity studies.

## 1. INTRODUCTION

Modularity is a widespread concept in modern science that emerged from the need to parcellate large, complex systems into smaller, hierarchically nested components (Simon, 1962). The study of modularity is commonplace in all biological disciplines because modularity affects the way complex biological systems, from genomes to ecosystems, originate, function, and evolve (Schlosser and Wagner, 2004; Callebaut and Rasskin-Gutman, 2005; Wagner et al., 2007). In morphology, the study of modularity focuses mostly on identifying regions of the body with a coordinated change of shape, by measuring traits covariation using distance-based morphometrics or landmark-based geometric morphometrics (reviewed in Esteve-Altava, 2016). The historical origin of this approach traces back to the influential book *Morphological Integration* by Everett Olson and Robert Miller (1958) in the context of zoological studies, and to the seminal paper on *The ecological significance of correlation pleiades* by Raissa Berg (1960) in botanical studies. Morphological modules identified on the basis of shape variation belong to the category of variational modules (Wagner and Altenberg, 1996; Eble, 2005; Wagner et al., 2007). Variational modularity has been the focus of many scholarly reviews in recent years (e.g., Klingenberg, 2008, 2010, 2014; Melo et al., 2016). In short, a variational module is a group of traits that vary coordinately (i.e., they are morphologically integrated *sensu* Olson and Miller) and, to some extent, they vary independently of other groups of traits. Up to two thirds of research studies on morphological integration and modularity analyze shape variation (Esteve-Altava, 2016), using variational module (more or less explicitly) as a synonym of morphological module. For this reason, in this essay we used variational modularity to refer to shape-variational modules as derived from morphometric analyses.

Morphological modules are also accessible to study on the basis of the topological interactions established among the constituent anatomical parts of a morphological system. This new conceptual framework uses network models and community detection algorithms to identify modules (Esteve-Altava et al., 2011; Rasskin-Gutman and Esteve-Altava, 2014). A network is a mathematical object that comprises two sets of elements: a set of nodes that represent the constituent parts of the system, and a set of links that connect pairs of nodes and represent interactions among these parts. Morphological systems are easily modeled as networks of body parts; in fact, networks are widely used already in neuroanatomy, where the brain is modeled as a network in which nodes represent neurons (or brain regions) and links represent synaptic connections or co-activation patterns (e.g., from fMRIs). Anatomical networks are abstract representations of an organism’s topology (Fig. 1): understanding topology as the way in which constituent parts are interrelated or arranged in the body (Rasskin-Gutman and Esteve-Altava, 2014). Although this quantitative approach is relatively new, a more general use of topology in morphology dates back to the beginnings of comparative anatomy and to Geoffroy Saint-Hilaire’s *principle of connections*. Ever since, connections among anatomical parts have been used in building theoretical models of morphological organization (e.g., Woodger’s *axiomatic method,* Rashevsky’s *bio-topological mapping,* and Riedl’s *diagrammatic morphotype*) and as a tool to establish homology between two body parts (see Rasskin-Gutman and Esteve-Altava, 2014 for an historical review). Because anatomical networks focus on explicit structural relations among body parts within an organism, independently of their variation, modules identified using anatomical network analysis belong to the category of organizational modules (Eble, 2005). An organizational module is a group of elements that establish more and/or stronger interactions within the group than outside it. Thus, the emphasize is placed on interactions among component parts, as an important constructional or functional property of form, whether interactions are defined based on structure (topology), pleiotropy, development, or performance (see Eble, 2005). Henceforth we used organizational modularity to refer to topology-organizational modules as derived from anatomical network analyses.

**Figure 1.**
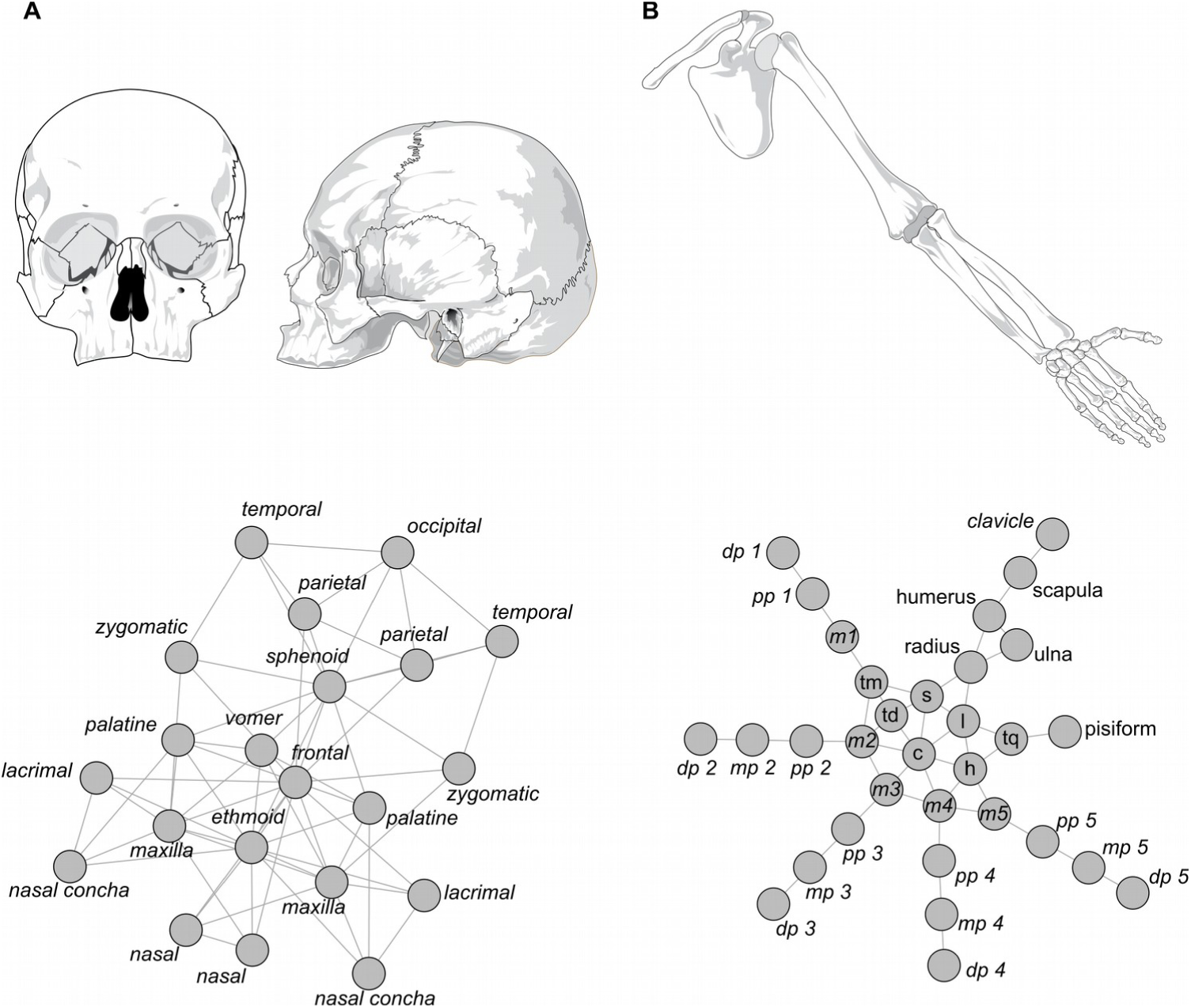
Network models of the skull (**A**) and of the upper limb (**B**). The first feature that catches the eye is the different organization of each network. On the one hand, the skull network shows a mesh-like organization, whereas the upper limb network shows a more star-like organization with serially connected nodes radiating from a central, small mesh. On the other hand, the skull network has a higher density of links (*K*_median_ = 5,*K*_min_ = 4, *K*_max_ = 13) than the upper limb network (*K*_median_ = 2, *K*_min_ = 1, *K*_max_ = 7). Networks are plotted using the Kamada-Kawai force-directed algorithm, which renders a natural layout for anatomical networks. Notice that the visual representation of a network model is trivial as long as the connections among nodes do not change.

Studies on variational and organizational modules have ontological and epistemological differences, although both approaches seek to parcellate complex morphological systems into highly integrated regions (Table 1). The source of these differences are (1) that each approach uses its own definition of form, and consequently, (2) that they use different methods to analyze organismal forms. Form is a rich concept that agglutinates information about proportion (i.e., size and shape) and structure (i.e., topology and arrangement), as well as other information related to the relative orientation and functional articulation of parts (Rasskin-Gutman and Buscalioni, 2001; Rasskin-Gutman, 2003). In this context of multiple layers of morphological information, variational modules deal with form at the level of proportions, while organizational modules deal with form at the level of structure. As a consequence, each approach uses a different set of proxies and formalisms. The raw data in morphometric-based variational modules are morphometric traits, such as linear distances and landmark coordinates, and the resulting mathematical objects analyzed are correlation or covariation matrices. In contrast, the raw data in network-based organizational modules are individual body parts and their topological relations, and the resulting mathematical objects are network models.

**Table 1.** Ontological and epistemological differences between variational and organizational modules.

|  | Variational Modules | Organizational Modules |
| --- | --- | --- |
| **Natural object** | complex morphological systems |  |
| **Level of study**<br>(*what is form?*) | proportions (*shape and size*) | structure (*topological arrangement*) |
| **Component parts** | morphometric traits | individualized anatomical parts |
| **Relation among parts** | correlation or covariation | topological boundaries |
| **Mathematical object** | correlation matrix | network model |
| **Module definition** | a group of traits that co-vary | a group of parts densely connected |
| **Identification**<br>(*exploration step #1*) | cluster analysis | community detection algorithm |
| **Validation**<br>(*exploration step #1*) | statistical test | optimization function $Q$<br>Wilcoxon rank-sum test |
| **Confirmation**<br>(*a priori hypothesis test*) | RV, CR, PLS | similarity |
| **Comparison**<br>(*with alternative partitions*) | EMMLi, z-score | similarity |

Identifying morphological modules (variational and organizational) from the empirical data described above, without any previous hypothesis of what are the actual modules, requires to use the adequate mathematical tools. In morphometrics we identify variational modules from the matrix of traits correlation using, for example, hierarchical clustering (Goswami, 2006) or graph modeling (Magwene, 2001). Note that in the latter method, graphs (or networks) are only used to summarize, or to help visualizing, statistical relationships among traits calculated from the correlation matrix; thus, the identification of modules does not rely on a network-based method as its name would suggest. However, it is possible to identify variational modules from correlation matrices using network-based methods (e.g., Perez et al., 2009; Suzuki, 2013; but see MacMahon and Garlaschelli, 2015 for methodological considerations). Moreover, correlation matrices can be constructed using a coordinate-based approach, treating each coordinate as a unit of variation (e.g., Klingenberg, 2008), or using a vector-based approach, treating each landmark (2D or 3D) as a unit of variation (e.g., Goswami and Polly, 2010; Goswami and Finarelli, 2016). Finally, we can validate the identified variational modules of the morphometric data using statistical tests (e.g., Fisher’s z-transformation and Student’s t-test, as in Goswami, 2006). On the other hand, in network-based methods we identify organizational modules using community detection algorithms. Broadly speaking, a network module is a group of nodes with more interactions (i.e., links) within the group than outside it. The identification of modules in networks has grown in sophistication in parallel with the application of networks to telecommunications, sociology, and biology (e.g., Palla et al., 2005; Newman, 2006; Fortunato, 2010); and within biology, most notably to ecology (e.g., Olesen et al., 2007), neurobiology (e.g., Sporns, 2011), and molecular biology (e.g., Guimerà and Nunes Amaral, 2005). Community detection algorithms seek to delimit modules using the topological information represented in the network model. However, identifying modules is computationally costly because of the large number of alternative partitions in which we can group the nodes of a network. Even in relatively small networks, such as the 21-node network of the human skull (described in Esteve-Altava et al., 2013) there are about 4.75 *x*10^14^ possible partitions. This is because the number of potential partitions of a network grows exponentially with the number of nodes, following the Bell’s numbers progression (Bell, 1938). There are many different algorithms to identify modules in networks, which vary in their heuristic approach. For example, some algorithms search the space of possible partitions by optimizing a quality function, while others use statistical inference on generative models, dynamic diffusion or spin processes (reviewed in Fortunato, 2010). Common validation methods include (1) the quantification of a function that measures the overall quality of the partition (which is usually the same used in optimization methods, see Eq. 1 below) according to the observed *vs*. expected links within and between modules (Newman and Girvan, 2004), (2) evaluating every module individually to meet a mathematical definition of module (Fortunato and Barthelemy, 2007), or (3) calculating the significance of modules using statistical tests (e.g., a Wilcoxon rank-sum test on internal *vs*. external number of links) or bootstrapping. Unfortunately, identifying network modules and validating network partitions are still open problems without universal agreed solutions (Fortunato and Hric, 2016).

Processes taking place at genetic, developmental, and functional levels, and across ontogenetic and evolutionary scales, are causally related to the emergence of morphological modularity (review in Klingenberg, 2008, 2014; Melo et al., 2016). Thus, a fairly common experiment consists in testing whether an *a priori* hypothesis of modularity based on information from one or more of these levels matches the morphological modules observed empirically in ontogeny or evolution (Esteve-Altava, 2016). Testing the fit of variational modules (or more generally, of traits covariation) to genetic, developmental, and functional hypotheses has a long tradition in morphology (see, e.g., Cheverud, 1982, 1989, 1996; Zelditch, 1988; Zelditch and Carmichael, 1989). There are various methods available to carry out such confirmatory tests, of which the most popular one in recent times is the Escoufier’s RV coefficient (Klingenberg, 2009). However, some authors have raised concerns about the reliability of RV coefficients and proposed alternative methods to validate *a priori* hypotheses of variational modularity. For example, Garcia, Oliveira, and Marroig have proposed the modularity hypothesis index (MHI, Garcia et al., 2015), which renders lower type I and II error rates than the RV; Adams has proposed the covariance ratio (CR, Adams, 2016), which (unlike RV) is not sensible to the size of the sample and to the number of variables, thus, allowing to perform comparisons across different data sets; lastly, Goswami and Finarelli have proposed an approach based on maximum likelihood and the Akaike information index to select among alternative hypotheses of modularity (EMMLi, Goswami and Finarelli, 2016). Note that this latter method would allow to compare competing partitions, such as those previously validated by one or more of the former methods. On the other hand, under the network-based approach, testing the fit of organizational modules to *a priori* hypotheses of modularity rely on measuring the similarity between two alternative partitions. These methods include measures based on pair counting, cluster matching, and information theory (Fortunato, 2010; Fortunato and Hric, 2016), all of which estimate to what extent the partition identified on a topological basis resembles a previously known partition based on metadata (e.g., genetic, developmental, and/or functional modules) or another algorithm. For example, in the context of morphological organizational modules, the normalized mutual information index (NMI, Danon et al., 2005) has been used to measure the similarity between the modules identified in the networks of the human limbs and various hypotheses based on the function and developmental origin of bones and muscles (Diogo et al., 2015).

Even though variational and organizational modularity differ in their epistemological and ontological basis, both approaches face similar challenges: the identification of reliable modules, their validation, and their comparison to alternative or *a priori* hypotheses. These challenges have been reviewed recently in the context of variational modularity and shape analysis (e.g., Goswami et al., 2014; Klingenberg, 2014; Garcia et al., 2015; Adams, 2016; Adams and Collyer, 2016; Goswami and Finarelli, 2016; Melo et al., 2016) and we will not discuss them further. Here we focus on these challenges in the context of organizational modularity and anatomical network analysis, by presenting a working example of how to identify, validate, and compare network modules in the anatomical networks of the skull and the upper limb of humans. Then, we discuss some ideas about how to integrate variational and organizational approaches. Although, it is not well-known whether, and how, variation and organization work together in structuring and shaping the form of organisms (but see, e.g., Perez et al., 2009; Esteve-Altava et al., 2013; Suzuki, 2013), the hope is that by bridging the gap between them we will have a better understanding of morphological modularity, and possibly help to tackle challenges on both sides.

## 2. STUDYING ORGANIZATIONAL MODULES USING NETWORK ANALYSIS

This section summarizes the process of identifying, validating, and comparing organizational modules using community detection algorithms and related methods. As an example, we used the anatomical networks of the skull and of the upper limb skeleton of humans. First, we introduce the concept of network model and how it formalizes the organization of morphological parts. Then, we present alternative approaches to evaluate the quality of partitions and of individual modules, which we apply afterward to validate the modules identified using a classic community detection algorithm based on the structural equivalence or topological overlap of nodes (GTOM, Ravasz et al., 2002). As we will see, the results of this approach will highlight most of the challenges we face when studying modularity in anatomical networks. To tackle these challenges, we also explored the use of a more sophisticated community detection algorithm based on local optimization of statistically significant communities (OSLOM, Lancichinetti et al., 2011). We close this working example by quantifying the similarity between network partitions and alternative partitions based on biological criteria, using information theory measurements. For the most part of the analysis we have used the free software *R 3.3.1* and the package *igraph 1.0.1*, unless otherwise stated; the source code and the network models are available as *Supplementary Materials*. The software to run OSLOM is available from the author’s page (www.oslom.org).

### 2.1. Anatomical Network Modeling

An anatomical network formalizes the way in which body parts are topological related, and, as such, it is a model of the organization of a morphological structure. Topological relations (connections) also embody developmental and functional interactions that take place between two body parts. For example, connections among skull bones are primary sites of bone growth and remodeling, while connections among limb bones are mobile articulations. Figure 1 shows the network models of the human skull (first published in Esteve-Altava et al., 2011) and of the human upper limb (first published in Diogo et al., 2015), in which nodes represent bones and links represent their physical joints (i.e., in the skull, craniofacial sutures and synchondroses; in the limb, mainly synovial joints).

A graph is a mathematical object that comprises a set of elements (vertices) and a set of pairwise-relations among elements (edges). A network is a graph with a non-trivial topology (e.g., not regular or random), although most often the terms graph and network are used as synonyms. Likewise, nodes (*N*) and links (*K*) are used as synonyms of vertices and edges, respectively. For simplicity, we modeled the networks of our examples as undirected (i.e., links have no direction) and unweighted (i.e., links are either present or absent), which is the simplest type of network. A network can be mathematically formalized as a binary adjacency matrix (*A_ij_*) of dimension *N* x *N*, in which the presence of a link between nodes *i* and *j* is coded as 1 and the absence as 0.

### 2.2. Definition of Module and Validation Partitions

A module (i.e., a community in network theory) is a subset of nodes more strongly connected with each other than with nodes outside the subset. To estimate how well a given partition of the network identifies the modules, Newman and Girvan (2004) defined the parameter *modularity* (commonly referred as *Q*),

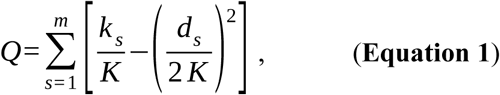

where *m* is the number of modules of the partition, *k_s_* is the number of links within module *s*, *d_s_* is the total number of links of nodes in *s* (both inside and outside *s*), and *K* is the total number of links in the network.

The parameter *Q* quantifies how strongly connected are the modules identified compared to a randomization of the network. *Q* is 0 when the number of links within modules is no better than expected in the randomization; higher values indicate a stronger modularity than expected, being *Q* = 1 the theoretical maximum. In practice, Newman and Girvan reported that strongly modular networks show values between 0.3 and 0.7. The expected error of *Q* can be calculated using a jackknife procedure where each link is considered as an independent observation. It is worth noticing that the value of *Q* is specific of each partition and network; thus, we can use it to compare among different partitions of a same network, but not to compare two different networks. In short, one network is not *more modular* than another because it has a higher value of *Q* for its best partition.

Equation 1 also includes the condition that one group of nodes has to fulfill to be a module, that is, having relatively more connections within the module than outside, which corresponds with the definition of module:

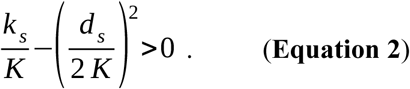

As a consequence, it is possible to have a partition of a network in which not all groups of nodes are modules according to Eq. 2. In turn, individual modules fulfilling Eq. 2 might have, in turn, sub-modules that also fulfill Eq. 2, but where not identified by the community detection algorithm. This situation produces a resolution limit in those community algorithms that directly or indirectly seek to find the partition of the network that renders the maximum modularity (Fortunato and Barthelemy, 2007). The underlying reason of this resolution limit is precisely that most networks have a hierarchical grouping of nodes into nested sub-modules. Alternatively, we can also validate each module of a partition statistically, for example, using a Wilcoxon rank-sum test on internal (*k_s_*) *vs*. external (*d_s_* – *k_s_*) number of links. Here we test the null hypothesis that there is no statistical difference between the number of internal and external links against the alternative hypothesis that the number of internal links is greater than the number of external links (i.e., the definition of a module). In our example, we used the Wilcoxon rank-sum on the modules identified by the first of the community detection algorithms used.

### 2.3. Identifying Modules with Community Detection Algorithms

The first community detection algorithm shown is based on a hierarchical clustering of the generalized topological overlap similarity matrix among nodes (GTOM, Ravasz et al., 2002), which is a classic method that uses a heuristic approach to identify modules. Heuristic methods are designed to overcome the otherwise computationally costly task of seeking and evaluating all the possible partitions of the network, by using an *a priori* reasoning of which nodes we would expect to group together. The heuristic of GTOM is that nodes that connect to the same other nodes (i.e., share neighbors) have a higher chance to belong to a same module.

The topological overlap between two nodes is the number of common neighbors between two nodes, defined as

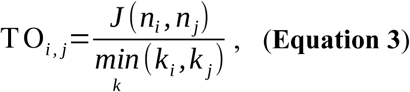

where *J*(*n_i_*,*n_j_*) is the number of neighbors in common between nodes *i* and *j*. *TO* is 1 when the two nodes share all their neighbors, that is, they connect to exactly the same other nodes. *TO* is 0 when the two nodes have no neighbor in common. By calculating the topological overlap over all pairs of nodes we get the GTOM, which is equivalent to a distance or dissimilarity matrix.

We can group nodes into clusters by using an agglomerative hierarchical cluster analysis on GTOM (in our example we used the average-linkage as in Ravasz et al., 2002; Esteve-Altava et al., 2013). The output is a hierarchical grouping of nodes, as in the two dendrograms shown in Figure 2. In order to identify the modules of the network we need then to decide at what level to cut the dendrogram. To make this decision we used (as it is customary in most hierarchical algorithms) the parameter *Q* explained before (Eq. 1). Thus, we measured *Q* for each possible partition of the dendrogram to identify the *best* partition, which is the one having the highest *Q* or *Q_max_* (Fig. 2; perpendicular dashed line in *red*). We can then calculate the statistical significance of each module or, as it is the case, of any cluster of the dendrogram, to evaluate the quality of each individual module identified by cutting the dendrogram at the level of *Q*_max_ (Fig. 2; circles on the dendrogram clusters).

**Figure 2.**
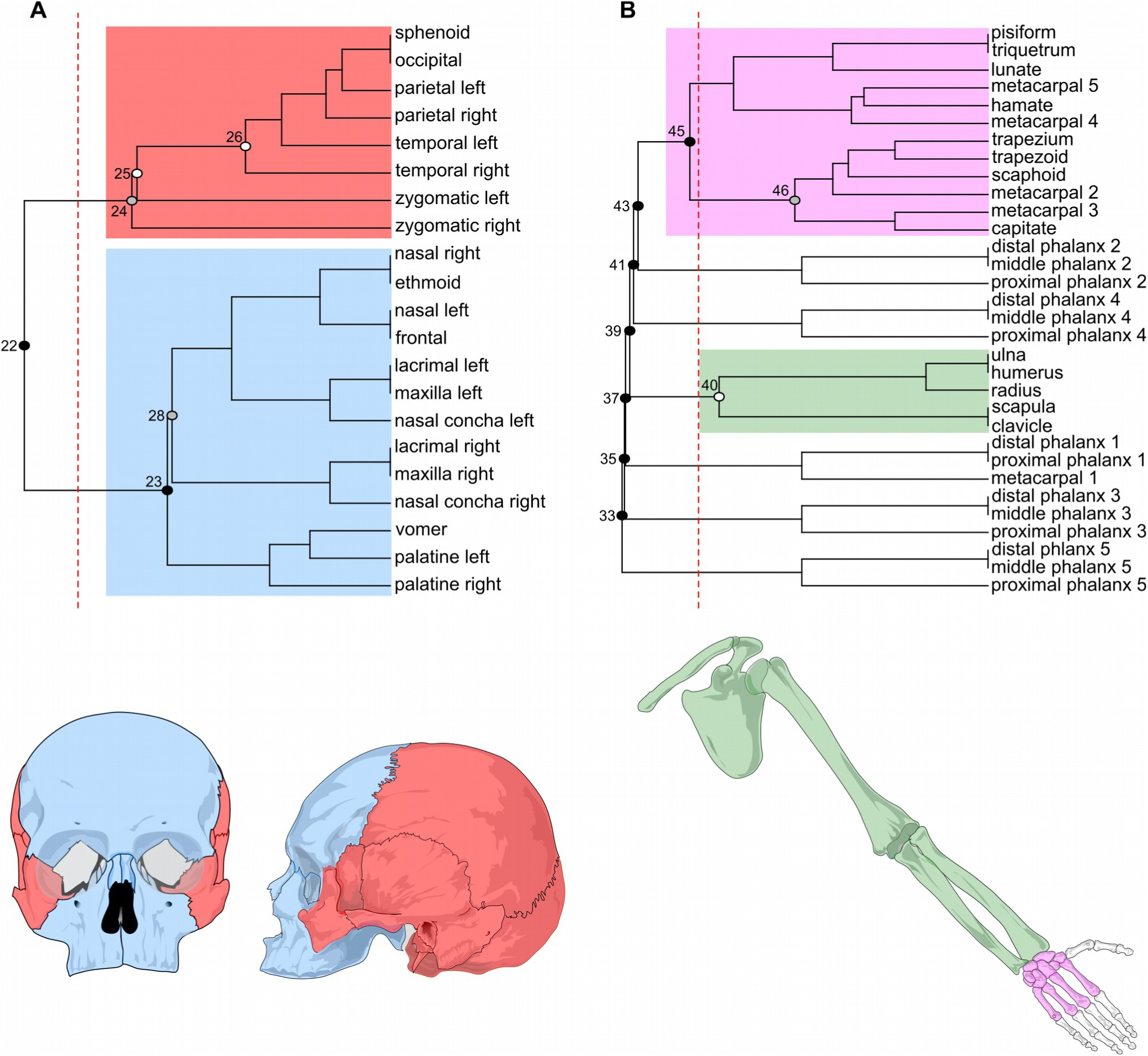
Modules identified with GTOM in the human skull (**A**) and upper limb networks (**B**). The dendrogram grouping the bones comes from the hierarchical cluster analysis of the TOM. The perpendicular dashed line in red indicates the partition of the dendrogram having the highest modularity (skull: *Q*_max_ = 0.27, *Q_error_* = 0.05; upper limb: *Q*_max_ = 0.52, *Q_error_* = 0.08). The circles on each bifurcation of the dendrogram indicate the statistical significance of that cluster in a Wilcoxon rank-sum test of internal *vs*. external connections: *black*, p-value < 0.001; *gray*, p-value < 0.01; *white*, p-value < 0.5; bifurcations without a circle are non-significant.

Using the GTOM algorithm the human skull network shows a relatively weak (*Q*_max_ < 0.3) modular partition in two modules. One module groups mainly the bones of the neurocranium (cranial base and vault) and the other groups the bones of the face (Fig. 2A, in *red* and *blue*, respectively). Both modules (clusters 24 and 23 in the dendrogram) are statistically significant, and include some sub-modules that are also significant (clusters 26 and 28) together with non-significant groups (e.g., vomer-palatines cluster) and singletons (e.g., the zygomatics). This pattern suggest that the human skull might have a partially hierarchical modularity, with some nodes having a lesser contribution. In contrast, the upper limb network shows a strong modular partition (*Q*_max_ > 0.5) in eight modules (Fig. 2B). One module groups the bones of the pectoral girdle, stylopod, and zeugopod (in *green*); two modules group the bones of the wrist (in *purple*); and five modules group the phalanges of each of the five digits. However, only two of these modules are statistically significant (clusters 40 and 46), which means that the GTOM algorithm returns *bad* modules as part of the best partition. Thus, we could ask whether is possible to find higher in the hierarchy a significant module without non-significant sister modules, so that all modules identified are significant. For example, in Figure 2, the *purple* module corresponds to cluster 45, which include the significant cluster 46 plus its sister cluster that is not significant. This shows that in a partition of the network using the *Q*_max_ a significant module can split into two sub-modules, not all of which being significant. In fact, the only partition of the upper limb network where all its modules are significant is the one-module partition (cluster 33), which would indicate that the whole network cannot be further divided into statistically significant modules (i.e., it is not modular, but fully integrated).

These examples illustrate two of the difficulties that the identification of modules in anatomical networks face: (1) the identification of weak partitions (as in the skull) and (2) of non-significant modules (as in the upper limb), which may be related to each other. The first difficulty is inherent to small size networks (e.g., tetrapod skull networks have between 20 and 60 nodes, Esteve-Altava and Rasskin-Gutman, 2014). The small size of networks hinders a correct statistical evaluation of their modules, in particular, when modules comprise only a few nodes (e.g., half of the modules of the upper limb had only three nodes); a small size also makes more difficult to discriminate between order and stochasticity in the connectivity patterns of the whole network. The second difficulty is imposed by the algorithm we choose. Many algorithms deal with the identification of rather simplistic modular organizations, letting aside (or underestimating) the presence of nested or overlapping modules, or even a partial or total lack of modularity. For example, it is possible that the anatomical networks of our examples are not truly hierarchical at this level, or that there is some degree of overlapping between their modules, or that these networks are not modular in part or in its wholeness after all.

To tackle these difficulties we used a second community detection algorithm based on the local optimization of statistically significant communities (OSLOM, Lancichinetti et al., 2011). OSLOM is specifically designed to identify significant modules locally, as well as the presence of hierarchical organization, overlapping modules (i.e., covers), partial modularity, and singletons (i.e., nodes not assigned to any module). Here, the module’s significance is taken as a fitness function that measures the probability of that module in a network without modularity (i.e., a randomization of the empirical network that keeps the same degree distribution). This probability is returned for every module identified as an estimation of the probability to find this module in the randomized network (bs). In short, the algorithm optimizes the module’s significance by iteratively adding and deleting nodes, looking for the most significant configuration available. This process is then iterated at a higher level to look for hierarchical groups. Because OSLOM evaluates the significance of modules individually, it can recover overlapping modules. Moreover, because the algorithm focuses on how individual nodes rise or lower the local significance of modules, it can also identify nodes (or groups of nodes) that do not fit within any module (i.e., singletons).

In contrast to the first community detection algorithm presented, OSLOM is stochastic, which means that the output results may vary from one run to another. OSLOM returns the results of the majority consensus, that is, the result found in more than 50% of the runs. Finally, two parameters need to be specified explicitly: the *tolerance*, which controls the significance threshold of modules; and the *coverage*, which controls whether to merge or to split sub-modules; together they affect the number of modules identified and their size. The authors advice that if the network lacks of a well-defined modularity, the choice of parameters values might affect the results. Thus, for each anatomical network we ran 1000 iterations, setting the coverage to 0.5 (the default value), and testing tolerances between 0.1 (default) and 0.5.

OSLOM returns two overlapping modules in the human skull network and one module plus a group of singletons in the upper limb network (Figure 3). For the skull network, OSLOM returns a slightly different arrange of modules depending on the value of tolerance (Fig. 4). For tolerance = 0.1, it returns no modules, which indicates that the network is highly integrated. For tolerance = 0.11, it identifies a *core-cranial module* that includes the occipital, sphenoid, parietal, and temporal bones (*bs* = 0.017). For larger values of tolerance (between 0.12 and 0.2), it identifies two modules that are similar to the cranial and facial modules identified by the first algorithm. In all instances, the cranial and the facial modules overlap in the frontal and zygomatic bones, which are shared between the modules. Interestingly, in most cases only one of the two zygomatic bones (left or right) participates in the overlap, which one does it specifically varies from rune to run, but since both have equally connected to both modules this difference is trivial. For this reason we consider both zygomatic bones as part of the overlap between the modules (Fig. 3A). Sporadically, for tolerance = 0.2, a broader overlap occurs which also includes the sphenoid bone (Fig. 4; *blue circle*). In general, the cranial module has a better estimated posterior significance that the facial module for all tolerance values, which means that a module like the cranial one is less likely to occur in a randomized network. In contrast, for the upper limb network, OSLOM returns only one module *(bs* = 0.153) grouping together the bones of the girdle, the stylopod (humerus), and the zeugopod (radius and ulna); while all the bones of the autopod (wrist and fingers) are not assigned to any module (i.e., they are singletons, see above). The one module identified corresponds to the statistically significant module identified using the GTOM algorithm (Fig. 2B; cluster 40, in *green*); the other non-significant modules identified using GTOM are not returned by OSLOM, where these bones are singletons (Fig. 3B, in *gray*).

**Figure 3.**
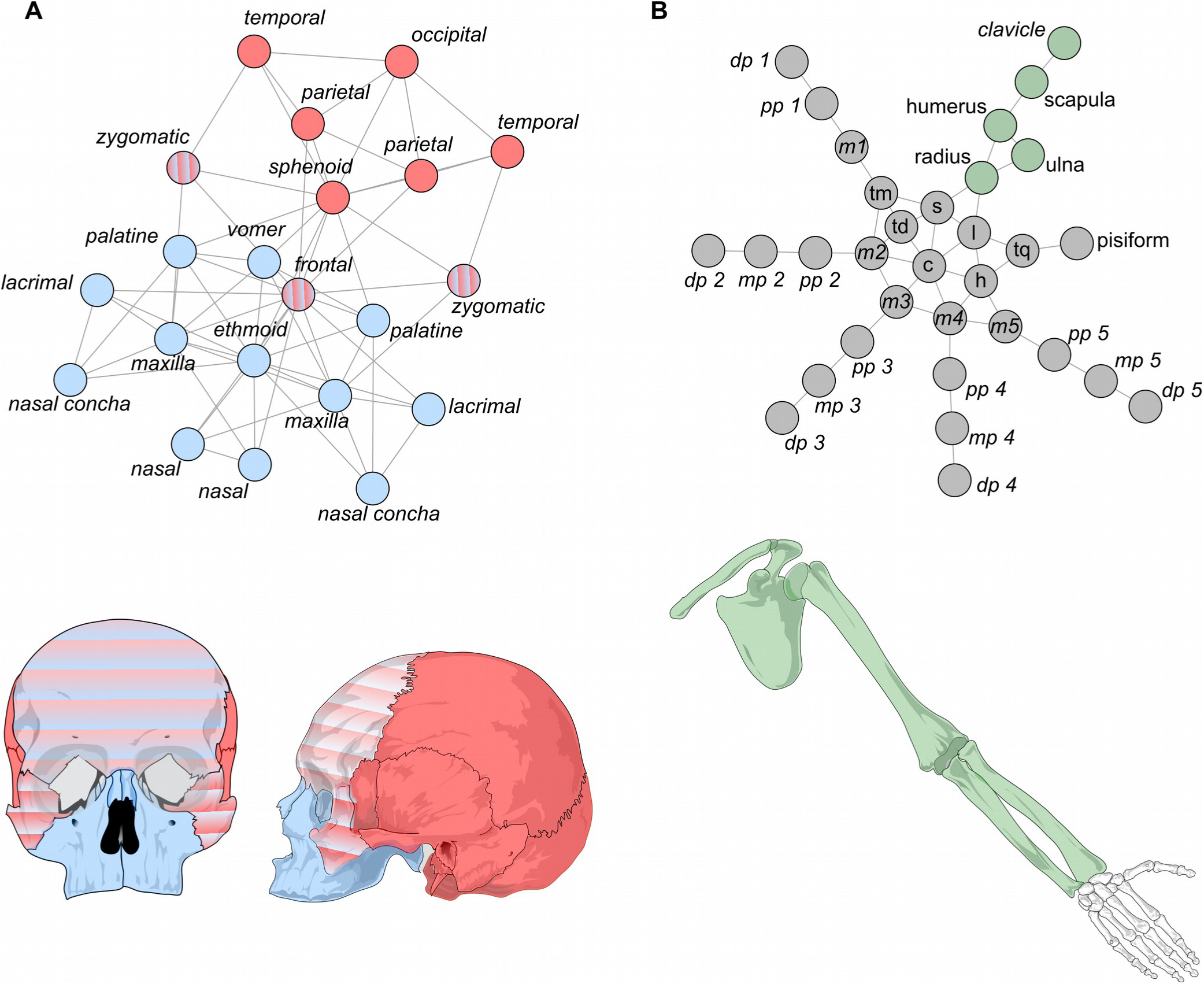
Modules identified with OSLOM in the human skull (**A**) and upper limb networks (**B**). For the skull, two modules were identified, one cranial (*red*) and one facial (*blue*), which overlap at the frontal and zygomatic bones (*red-blue gradient pattern*). For the upper limb, only one module was identified (*green*), while the rest of the bones do not form a module (*gray*).

**Figure 4.**
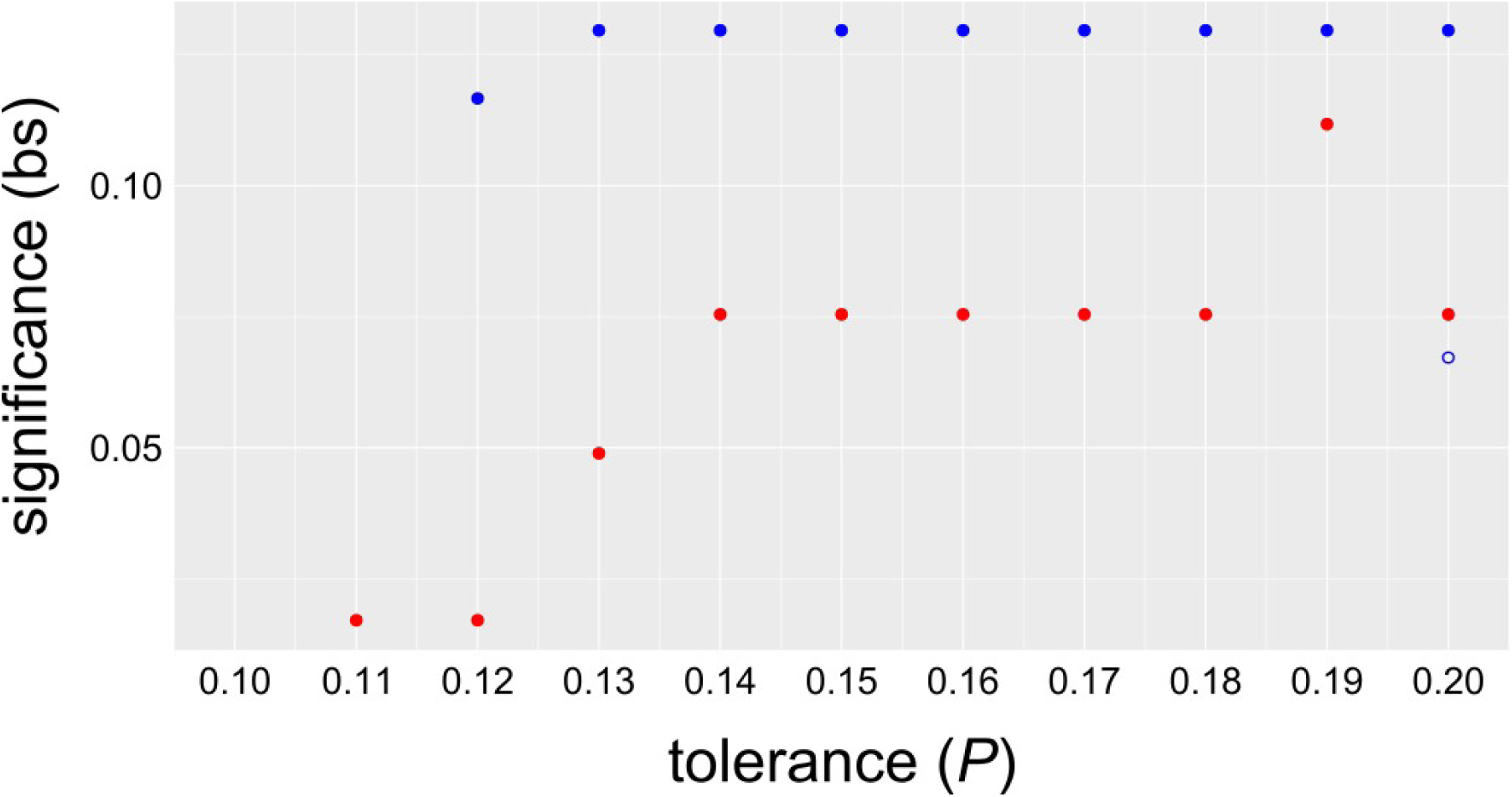
Significance of the cranial (*red dots*) and facial module (*blue dots*) identified with OSLOM for a range of tolerance values. The facial module is only identified for tolerances greater than 0.11 and always with a significance higher than the cranial module. The *blue circle* indicates the value of an alternative, less frequent facial module that includes also the sphenoid bone (see *Main Text*).

### 2.4. Comparing between two partitions (or *how to test a Ho*)

We compared the partition of the skull and upper limb networks identified with the two algorithms to alternative partitions based on different developmental criteria. For the skull (Table 2), we compared the partitions by GTOM and OSLOM to a partition of bones by their ossification mechanism (dermal, endochondral, and mixed) and to a partition of bones based on their cellular origin (mesoderm, neural crest, and mixed). For the upper limb (Table 3), we compared the partitions by GTOM and OSLOM to two partitions of the limb based on its developmental patterning, the traditional one (girdle, stylopod, zeugopod, and autopod) and a variant that also includes the mesopod region (girdle, stylopod, zeugopod, mesopod, and autopod). For simplicity, we considered all the singletons of the upper limb (i.e., bones not assigned to any module) as forming one module of their own.

**Table 2.** Divisions of the human skull compared.

| Bone | Ossification | Cell origin | GTOM | OSLOM |
| --- | --- | --- | --- | --- |
| *Occipital* | Mixed | Mesoderm | Cranial | Cranial |
| *Parietals* | Dermal | Mesoderm | Cranial | Cranial |
| *Temporals* | Mixed | Mixed | Cranial | Cranial |
| *Sphenoid* | Endochondral | Mixed | Cranial | Cranial |
| *Zygomatics* | Dermal | Neural crest | Cranial | Overlap |
| *Frontal* | Dermal | Neural crest | Facial | Overlap |
| *Ethmoidal* | Endochondral | Neural crest | Facial | Facial |
| *Nasals* | Dermal | Neural crest | Facial | Facial |
| *Maxillas* | Dermal | Neural crest | Facial | Facial |
| *Lacrimal*s | Dermal | Neural crest | Facial | Facial |
| *Palatines* | Dermal | Neural crest | Facial | Facial |
| *Nasal conchae* | Dermal | Neural crest | Facial | Facial |
| *Vomer* | Dermal | Neural crest | Facial | Facial |

**Table 3.** Divisions of the human upper limb compared.

| **Bone** | **Dev. pattern I** | **Dev. pattern II** | **GTOM** | **OSLOM** |
| --- | --- | --- | --- | --- |
| Clavicle | Girdle | Girdle | Cluster 40 | Module |
| Scapula | Girdle | Girdle | Cluster 40 | Module |
| Humerus | Stylopod | Stylopod | Cluster 40 | Module |
| Radius | Zeugopod | Zeugopod | Cluster 40 | Module |
| Ulna | Zeugopod | Zeugopod | Cluster 40 | Module |
| Trapezoid | Autopod | Mesopod | Cluster 46 | Singleton |
| Trapezium | Autopod | Mesopod | Cluster 46 | Singleton |
| Scaphoid | Autopod | Mesopod | Cluster 46 | Singleton |
| Lunate | Autopod | Mesopod | Cluster 45 (*ns* branch) | Singleton |
| Triquetrum | Autopod | Mesopod | Cluster 45 (*ns* branch) | Singleton |
| Pisiform | Autopod | Mesopod | Cluster 45 (*ns* branch) | Singleton |
| Hamate | Autopod | Mesopod | Cluster 45 (*ns* branch) | Singleton |
| Capitate | Autopod | Mesopod | Cluster 46 | Singleton |
| Metacarpal 1 | Autopod | Autopod | Digit 1 | Singleton |
| Metacarpal 2 | Autopod | Autopod | Cluster 46 | Singleton |
| Metacarpal 3 | Autopod | Autopod | Cluster 46 | Singleton |
| Metacarpal 4 | Autopod | Autopod | Cluster 45 (*ns* branch) | Singleton |
| Metacarpal 5 | Autopod | Autopod | Cluster 45 (*ns* branch) | Singleton |
| Proximal phalanx 1 | Autopod | Autopod | Digit 1 | Singleton |
| Distal phalanx 1 | Autopod | Autopod | Digit 1 | Singleton |
| Proximal phalanx 2 | Autopod | Autopod | Digit 2 | Singleton |
| Middle phalanx 2 | Autopod | Autopod | Digit 2 | Singleton |
| Distal phalanx 2 | Autopod | Autopod | Digit 2 | Singleton |
| Proximal phalanx 3 | Autopod | Autopod | Digit 3 | Singleton |
| Middle phalanx 3 | Autopod | Autopod | Digit 3 | Singleton |
| Distal phalanx 3 | Autopod | Autopod | Digit 3 | Singleton |
| Proximal phalanx 4 | Autopod | Autopod | Digit 4 | Singleton |
| Middle phalanx 4 | Autopod | Autopod | Digit 4 | Singleton |
| Distal phalanx 4 | Autopod | Autopod | Digit 4 | Singleton |
| Proximal phalanx 5 | Autopod | Autopod | Digit 5 | Singleton |
| Middle phalanx 5 | Autopod | Autopod | Digit 5 | Singleton |
| Distal phalanx 5 | Autopod | Autopod | Digit 5 | Singleton |

*ns*, non-significant

We compared partitions using an index based on information theory, the *normalized mutual information index* (NMI, Danon et al., 2005). NMI measures the similarity of two partitions based on the additional amount of information needed to infer one partition from the other (similar partitions would need less information) and normalizes it by dividing by the arithmetic mean of the entropy of both partitions as

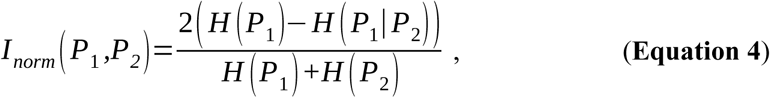

where *H*(*P_1_*) is the Shannon entropy of the first partition and *H*(*P_1_*|*P_2_*) is the conditional entropy of the first partition given the second partition. NMI is 1 when the two partitions are identical, and it is 0 when they are totally different. For convenience we express the similarity between to partitions in percentages.

The partitions of the skull based on the ossification mechanism and on the cellular origin of bones are different, 46.8% similarity, which is almost half of the similarity between the results of GTOM and OSLOM, 70.8%.

This result is expected because the two latter partitions are both based on topology (GTOM *vs.* OSLOM), whereas the two developmental partitions are based on different criteria (ossification *vs*. cell origin). Partitions by GTOM and OSLOM are more similar to that based on cellular origin of bones, 52.6% and 68.3%, respectively, than to that based on ossification mechanism of bones, 24.7% and 30.8%, respectively. In both comparisons, OSLOM outperforms GTOM in identifying a division of the human skull similar to those based on developmental criteria.

The partitions of the upper limb based on the alternative developmental patterning of the limb (with and without a mesopod) are similar, as we would expect, 70.8%, which is almost the double of the similarity between partitions of GTOM and OSLOM, 35.2%. It is surprising here the low similarity between both algorithms, which might be related with the number of modules identified by each algorithm, seven and one, respectively. The partition by GTOM is more similar to that of the developmental patterning with the mesopod, 45.1%, than without the mesopod, 33%; whereas the partition by OSLOM is more similar to that without the mesopod, 84%, than with the mesopod, 56.1%. In both cases, again, OSLOM outperforms GTOM in identifying a division of the human upper limb similar to those based on developmental patterning.

### 2.5. Biological Interpretation of Network-based Organizational Modules

What does it mean for a group of bones to be in a same network module? The answer to that question depends on what are the actual biological functions of the topological interactions or relations that we formalized as the network links. This is best illustrated by our example of the human skull, which consistently shows a modular partition in two modules, one grouping the bones of the cranial vault and base (cranial module) and one grouping the bones of the face (facial module). We built the network model of the human skull by formalizing craniofacial sutures and synchondroses as the links of the networks. Among the most important functions of the sutures and synchondroses of the skull is to act as primary sites of bone growth and remodeling (Opperman, 2000; Rice, 2008; Lieberman, 2011). In other words, a link represents a shared (i.e., correlated) growth of the two bones linked. Because a network module is a group of bones more densely connected among them than to other bones outside the module, bones that belong to the same module share more growth relations among them (on average) than with other bones. Thus, we would interpret the facial and cranial modules as semi-independent units of growth (Esteve-Altava et al., 2013). Alternative interpretations of network modules are possible because connections among anatomical parts rarely carry one single biological function. For example, in addition to being growth sites, we know that connections among skull bones have an also an important biomechanical role, being key actors in processes of stress diffusion and tension release (Rafferty et al., 2003; Moazen et al., 2009; Curtis et al., 2013). In this context, we would interpret the cranial and facial modules of the skull as semi-independent biomechanical units.

Our example of the upper limb network is useful to illustrate the *a posteriori* interpretation of modules as evolutionary units or as constrains to evolvability. In the upper limb, both algorithms identify a well-defined module comprising the bones of the girdle, stylopod, and zeugopod; but algorithms differ in how to group the bones of the autopod. GTOM groups them en 7 different modules, whereas OSLOM finds they are all singletons with not clear modular organization. The fact that most of the autopod modules identified by GTOM are not significant supports the result of OSLOM. In the upper limb network, links represent physical articulation, via cartilaginous joints, among bones. This pattern of articulation in the limb is highly conserved in evolution, and deviations of this connectivity pattern to accommodate functional adaptations of the upper limb (e.g., to run, burrow, flight, swim, etc) take place mostly at the autopod level (Lewis, 1989). In fact, a similar pattern of connections between the girdle, stylopod, and zeugopod bones is already present in Devonian tetrapodomorphs, which still lack of a well-defined autopod (Clack, 2009). Thus, we can interpret this module as a highly integrated evolutionary unit, which imposes a constraint to its evolvability. In contrast, the bones of the autopod do not group in a module, being free to vary semi-independently of the proximal module and to accommodate functional needs without disrupting more proximal structures. The fact that the autopod is an evolutionary novelty of tetrapods (albeit its developmental homologies with fin rays, e.g., see Nakamura et al., 2016) reinforces the idea of its semi-independent evolvability.

## 3. BRIDGING THE GAP BETWEEN VARIATIONAL AND ORGANIZATIONAL MODULARITY

Are shape-variational modules causally related to topology-organizational modules? The answer to that question depends on the existence of an actual relationship between shape and topology in the generation of organismal forms. Some sort of relation exist between shape and topology due to the fact that landmarks covariation is constrained by the topological contiguity of the body parts on which landmarks are are located (Chernoff and Magwene, 1999; Magwene, 2001, 2008; Klingenberg, 2009). This was first reported in a study on the factors determining individual bone shapes covariation in the human skull by Karl Pearson and Wu Dingliang, who found that contiguity (i.e., adjacency or connection) between two bones correlates with a covariation of shape between them (Pearson and Woo, 1935). This study showed that the adjacency of bones (i.e., a connection in a network model) is the most important factor, after symmetry, in explaining the co-variation in shape of two skull bones. Unfortunately, the correlation between topology and shape has not been the subject of further experimental studies since then. As a consequence, it is unknown whether this correlation comes from a one-way causation (from topology to shape or the other way around) or from a two-way causal relationship; furthermore, it is possible that this correlation is caused by a third factor acting on both features (e.g., growth), or even, it might be an artificial correlation due to flaws in the design of the experiment.

The simplest way to explore whether shape-variational modules match with topology-organizational modules is to use organizational modules as null hypothesis of shape variation, to be tested with morphometric methods – a task which easier said than done. This hypothesis assumes that organizational modules, as derived from a network analysis of body parts, act as a map of correlations or co-variations imposing structural constraints on shape. Additionally, we might use an exploratory morphometric analysis to group bones according to their shape correlation and then use a similarity test, as the one shown in the previous section, to validate the match of both partitions. In any case, a well-rounded confirmatory analysis would use both validation approaches, ideally, using independent datasets.

Finally, information on shape variation might be directly included in the construction of the anatomical network model, so it is taken into account when we perform the community detection. For example, shape covariation of two bones might be used to weigh their connection; thus, we would have a weighted anatomical network where each link represent a topological connection pondered by the actual shape covariation between the two bones. Since we only use the covariation of connected nodes, the resulting mathematical object would be different of a direct network of the matrix of correlations (as in Perez et al., 2009; Suzuki, 2013).

## 4. CONCLUDING REMARKS

Morphological systems have a multi-level modularity, which is not limited to the underlying modularity of their generative processes and their consequences on shape, but it is also manifested at a morphological level in the structural organization of body parts. Anatomical network models and their analysis using community detection algorithms offer a new, complementary set of tools to identify morphological modules, and study how they change in development and evolution. We face the challenge now to further develop these tools in morphology, revealing the causal connections between structure and shape in the origin and evolution of organismal forms.

## 5. ACKNOWLEDGMENTS

I am thankful to Julio Hoyos and Rui Diogo for inviting me to present at the symposium *“Major Challenges for Vertebrate Morphology, Evolution, and Development Symposium”*, held at the 11^th^ International Congress of Vertebrate Morphology (2016 Washington, DC), which inspired this essay. This project received funding from the European Union’s Horizon 2020 research and innovation programme under Marie Skłodowska-Curie grant agreement No 654155.

